# Structural basis for the coiled-coil architecture of human CtIP

**DOI:** 10.1101/2021.03.05.434060

**Authors:** C. R. Morton, N. J. Rzechorzek, J. D. Maman, M. Kuramochi, H. Sekiguchi, R. Rambo, Y. C. Sasaki, O. R. Davies, L. Pellegrini

**Author notes:** The Francis Crick Institute, London NW1 1AT, UK.

## Abstract

The DNA repair factor CtIP has a critical function in Double-Strand Break (DSB) repair by Homologous Recombination, promoting the assembly of the repair apparatus at DNA ends and participating in DNA-end resection. However, the molecular mechanisms of CtIP function in DSB repair remain unclear. Here we present an atomic model for the three-dimensional architecture of human CtIP, derived from a multi-disciplinary approach that includes X-ray crystallography, Small-angle X-ray Scattering (SAXS) and Diffracted X-ray Tracking (DXT). Our data show that CtIP adopts an extended dimer-of-dimers structure, in agreement with a role in bridging distant sites on chromosomal DNA during recombinational repair. The zinc-binding motif in CtIP’s N-terminus alters dynamically the coiled coil structure, with functional implications for the long-range interactions of CtIP with DNA. Our results provide a structural basis for the three-dimensional arrangement of chains in the CtIP tetramer, a key aspect of CtIP function in DNA DSB repair.

## Introduction

Damage to the chemical structure of DNA is a constant threat to the genetic stability of the cell and the faithful transmission of genetic information. In response to DNA damage, complex cellular responses have evolved to signal the presence of genotoxic lesions and activate the appropriate repair pathway^1,2^. Double-strand breaks (DSBs) in the DNA double helix represent a highly dangerous injury that – if left unrepaired – can cause chromosomal rearrangements and genomic instability, a major predisposing factor to development of cancer and other pathologies. DSBs in eukaryotic cells are normally repaired by two conserved repair mechanisms: Non-Homologous End Joining (NHEJ)^3^, which re-joins directly the broken DNA ends with occasional loss of information at the damaged site, and Homologous Recombination (HR)^4^, a high-fidelity mode of repair prevalent during and after DNA replication, when a sister chromatid copy is available as a template for repair.

The choice of DSB repair pathway is influenced by complex cellular mechanisms that depend on the preferred recruitment to the DNA ends of specialised protein factors, which then determine the mode of DSB repair^5–7^. The extent of DNA-end resection is a critical factor in deciding the DSB repair pathway. Resection is normally prevented by the NHEJ end-binding Ku protein and the 53BP1-Rif1-Shieldin complex, which act to channel repair towards NHEJ. During S-phase, the role of the BRCA1-BARD1 complex becomes predominant in promoting extensive resection of DNA ends, thus favouring repair by HR.

The CtIP (CtBP-Interacting Protein) / RBBP8 (Retinoblastoma-Binding Protein 8) was first identified as a transcriptional co-repressor, but accumulating evidence has since established its paramount importance in maintaining genomic stability. Thus, CtIP has a critical role in promoting HR, acting together with BRCA1 and the Mre11-Rad50-Nbs1 (MRN) complex to initiate DNA-end resection^8,9^. Furthermore, CtIP mediates the cell-cycle control of repair choice upon CDK phosphorylation at T847^10^, a post-translational modification (PTM) that is necessary for efficient resection and is shared with its distant yeast orthologue Sae2^11^. Although its mechanism of action remains poorly understood, current evidence indicates that one important function of CtIP/Sae2 is to activate the endonucleolytic cleavage of blocked DNA ends by the Mre11 nuclease at the start of the resection process^12,13^. In addition to its direct involvement in the process of enzymatic resection of DNA ends, CtIP is a hub for protein-protein interactions, coordinating recruitment of repair factors to damaged DNA^9^. CtIP is also extensively post-translationally modified, and these PTMs are important for its repair function^9^. Recent evidence further shows that CtIP’s role in maintaining genomic stability extends to stabilisation of stalled forks during DNA replication^14–16^.

Our mechanistic understanding of the function of CtIP in HR is limited by lack of structural information. Such information has proved difficult to acquire as CtIP appears to be intrinsically disordered over most of its sequence, with the exception of its alpha helical N-terminal region^17^. Amino acid conservation points to the presence of functionally important N- and C-terminal domains^18–20^ that have been shown to bind DNA and the MRN complex^20–24^. An important mechanistic advance came from the demonstration that CtIP exists in a tetrameric form, due to the presence of a short tetramerisation motif at its helical N-terminus^21^. Tetramerisation is important for CtIP’s repair function, as the single-point mutation L27E that abolishes tetramer formation yields a protein that is deficient in HR^21^. Indeed, conservation of the tetramerisation motif in the fission-yeast orthologue Ctp1^22^ reinforces the important functional role of this oligomeric state in DSB repair.

The crystal structure of the tetramerisation motif, together with the predicted presence of C-terminally juxtaposed coiled-coil segments, suggested an elongated structure for CtIP, based on the head-to-head association of two parallel coiled-coil dimers^21^. Such an architecture is supported by a recent characterisation of full-length human CtIP showing that the tetrameric protein forms a dumbbell shape, in agreement with the proposed mechanism of tetramerisation^25^. The functional implication of such structural arrangement is that CtIP might be able to bridge distinct DNA molecules, and to position its conserved functionally important C-terminal domains at distant DNA sites. Biochemical evidence for such bridging behaviour, as well as for formation of higher-order assemblies, has recently been obtained^26,27^.

The tetramerisation motif of human CtIP spans amino acids 18 to 32, whereas the region with strongly predicted alpha helical conformation extends to residue 145 (**Figure 1A, B**). Furthermore, embedded within the coiled-coil region is a zinc-binding motif of unknown structure, where zinc coordination is shared between two conserved cysteine ligands in each chain of the dimeric coiled coil. Thus, the nature of the three-dimensional architecture of the CtIP N-terminus and how it affects CtIP function remains to be determined. In this paper, we present evidence – drawing on multiple experimental techniques – that the conserved helical sequence of CtIP juxtaposed to its tetramerisation motif forms a parallel coiled coil, providing experimental support for the proposed dimer-of-dimers architecture. Furthermore, we show that the presence of the zinc-binding motif in the middle of the coil allows for the generation of distinct geometries of the CtIP N-terminus.

**Figure 1.**
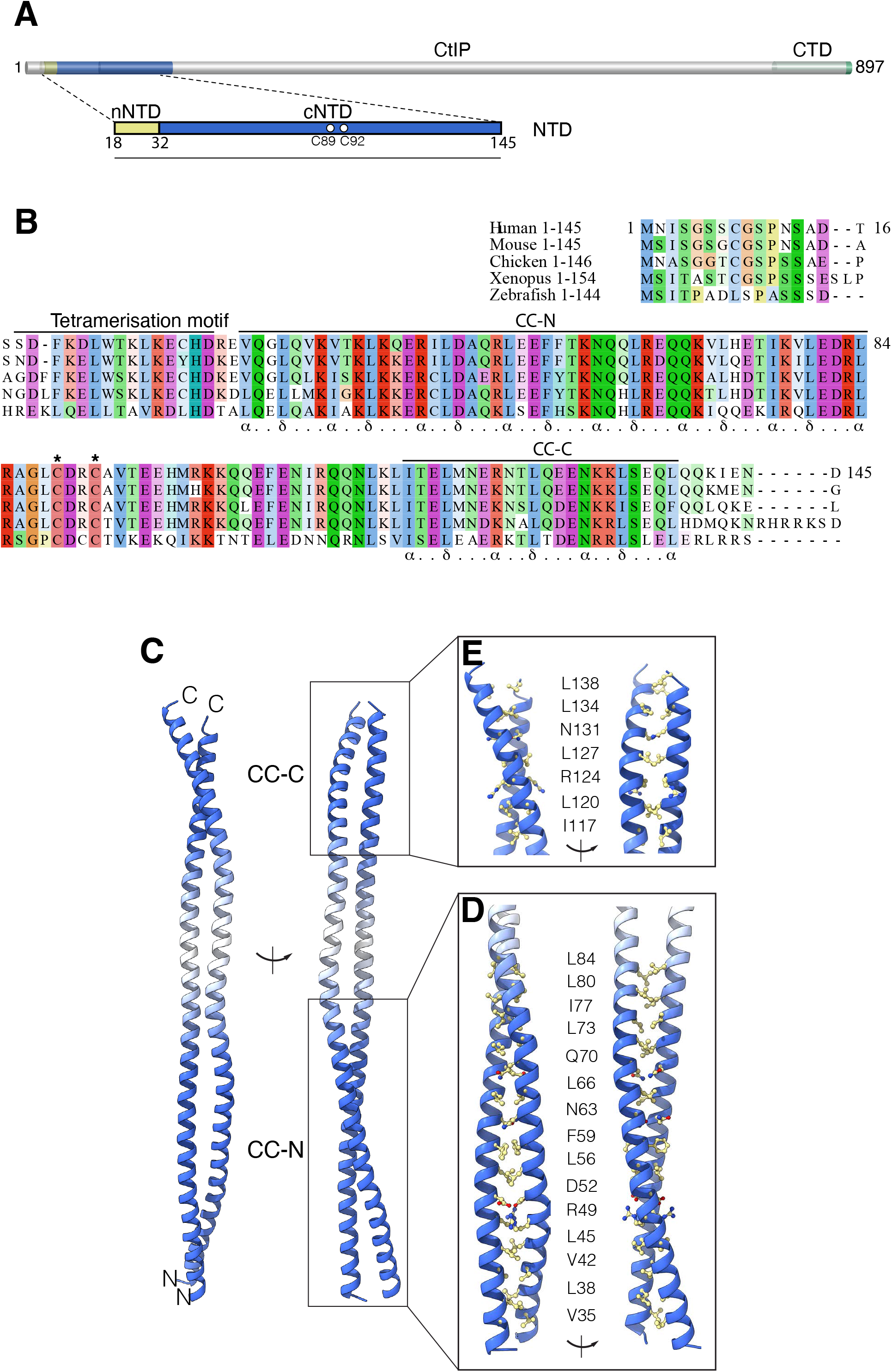
Coiled coil structure of CtIP-cNTD. **A** The domain structure of human CtIP. The amino-terminal domain of CtIP (NTD) is comprised of an N-terminal sequence (nNTD), responsible for CtIP tetramerisation, and a C-terminal coiled-coil sequence (cCTD) that includes a zinc-binding motif, with C89 and C92 as ligands. **B** Sequence conservation at the N-terminus of CtIP. The multiple sequence alignment is annotated to show the amino acid extent of CtIP’s tetramerisation motif and coiled coil CC-N and CC-C segments. **C** Ribbon drawing of the crystal structure of the CtIP-cNTD. Two views of the structure related by a 90° rotation around its coiled coil axis are shown. The coiled coil regions of the structure are coloured in blue, whereas the intervening helical sequence is coloured in white. The N- and C-terminal coiled coils are labelled CC-N and CC-C, respectively. **D** Structural details of the CC-N. The side chains of interdigitating residues in the coiled coil are shown as ball-and-stick models. Two rotated views are shown. **E** Same as in B, but for the CC-C.

## Results

### Crystal structure of the CtIP N-terminal coiled-coil

To improve our understanding of CtIP structure and function, we focused on the region that is juxtaposed to the C-terminus of the tetramerisation motif. In keeping with the nomenclature adopted for the CtIP N-terminal domain (NTD)^21^, we will refer to this sequence of CtIP as cNTD (C-terminal NTD), to distinguish it from the N-terminal part of the NTD (N-terminal NTD; nNTD) which comprises the tetramerisation motif (**Figure 1A, B**). Constructs designed for crystallisation started after leucine 27, which is critical for tetramerisation^21^, and extended to residue 145, therefore including the entire span of the predicted alpha helical region. Sparse matrix screening resulted in diffracting crystals for CtIP-cNTD construct 31-145; however, diffraction quality was poor and attempts to improve crystal quality were not successful.

The CtIP-cNTD binds zinc via cysteines C89 and C92^21^; the zinc-binding motif thus interrupts the extended helical sequence of the CtIP-NTD to generate two flanking coiled coil segments (**Fig. 1A, B**). Zinc binding is shared by the two chains of the CtIP-cNTD dimer (1 zinc : 2 cNTD stoichiometry), but it does not play a role in CtIP-cNTD dimerisation^21^. We reasoned that sub-stoichiometric incorporation of zinc in recombinant CtIP-cNTD might cause conformational heterogeneity preventing formation of well-ordered crystals. A double C89A, C92A mutant (Mut) CtIP-cNTD protein was prepared, which crystallised readily and showed improved diffraction properties. We determined its crystal structure at 2.8 Å resolution by molecular replacement, using as search template a composite homology model comprising the strongly predicted coiled-coil regions flanking the zinc-binding motif (**Supplementary table 1** and **Methods**).

The crystal structure revealed that the cNTD region spanning residues 31-136 folds in a highly elongated dimer of uninterrupted parallel alpha helices (**Figure 1C**). Within this helical structure, two coiled-coil regions – named here CC-N and CC-C – comprising amino acids V35 to L84 (CC-N) and I117 to L138 (CC-C), flank the zinc-binding site (**Figure 1E, D**). The coiled-coil structure of both CC-N and CC-C follows largely the expected heptad-repeat pattern of interdigitating hydrophobic residues. The residues surrounding the positions of cysteines 89 and 92, comprising the zinc-binding motif, interrupt the coiled-coil conformation and form instead two alpha helices running parallel to each other in the dimer (**Figure 1C**).

The presence of two cysteine ligands per CtIP chain and the observed 2:1 protein-to-zinc stoichiometry^21^ had led to the expectation of intermolecular coordination of one zinc ion by two CtIP chains, with the four cysteine ligands pointing inwardly towards the metal ion. Surprisingly, in the crystal structure the alanine residues occupying the position of the zinc-binding cysteines 89 and 92 point outwardly towards the solvent (**Figure 2A**), in a conformation that appears incompatible with shared coordination of a zinc ion (**Figure 2B**). The reason for this unexpected arrangement is unclear; it is possible that the presence of alpha helix-promoting alanine residues in the mutant protein had induced formation of a local helical conformation, leading to the observed continuous alpha helical structure for the entire CtIP-cNTD sequence.

**Figure 2.**
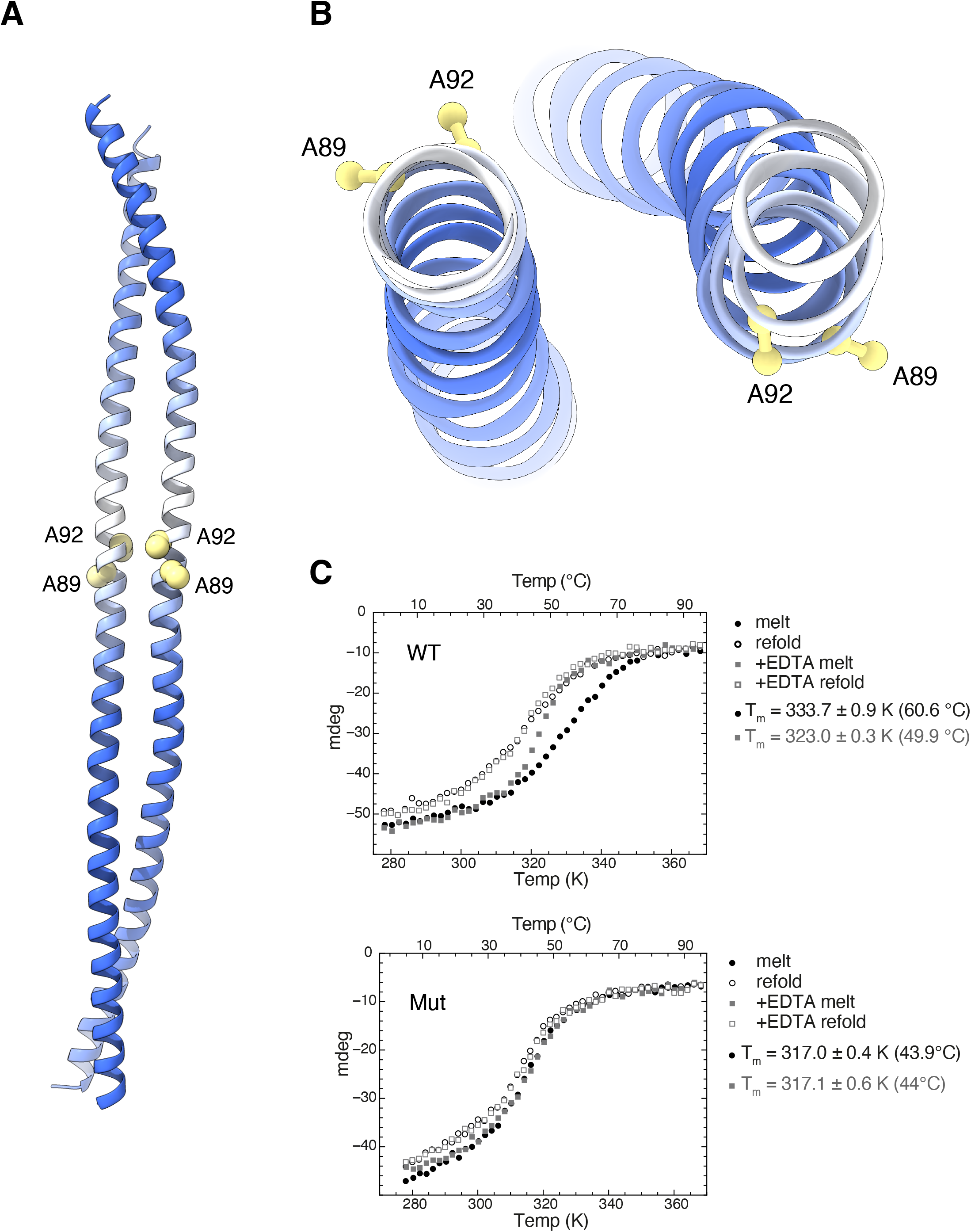
Characterisation of the cNTD mutant. **A** Conformational consequence of the double C89A, C92A mutation. The position of the alanine residues is shown by drawing their side chains as spacefill models, whereas the rest of the structure is drawn as in figure 1. **B** Top view of the cNTD structure, highlighting the outwardly pointing side chains of A89 and A92. **C** Melting and refolding CD curves of wild-type (top) and C89A, C92A mutant cNTD (bottom), in the presence and absence of EDTA.

### CD analysis of CtIP-cNTD

We decided to explore the conformation of the wild-type and Mut CtIP-cNTD proteins using circular dichroism (CD) melting experiments. The CD analysis shows that the wild-type protein is considerably more stable than the mutant (T_m, WT_ = 333.7 K *vs* T_m, Mut_ = 317.0 K) (**Figure 2C**), supporting a structural model where zinc crosslinks two CtIP chains, thus conferring additional stability to the coiled coil interactions of the protein dimer. In agreement with the model, treatment of the wild-type protein with the divalent ion-chelator EDTA reduced the T_m, WT_ (323.0 K) whereas it had no significant effect on the T_m, MUT_ (**Figure 2C**). Interestingly, the CD refolding curve of wild-type CtIP-cNTD did not coincide with its melt curve (**Figure 2C**), suggesting that its refolding is a complex process, as would be expected for a structure that contains a folded zinc-binding module in addition to helical structure. In contrast, the melt and refold curves of the mutant CtIP-cNTD protein overlap, indicative of a simpler conformational transition between folded and unfolded states.

### SAXS analysis of the CtIP-cNTD dimer

To investigate further the structure of CtIP-cNTD, we determined its size and shape by size-exclusion chromatography small-angle X-ray scattering (SEC-SAXS; referred to as SAXS here) (**Figure 3A** and **Supplementary table 2**). SAXS analysis revealed cross-sectional radii (*R*_*c*_) of wild-type and mutant proteins of 8.4 Å and 9.4 Å, respectively (**Supplementary Figure 1A, B**), within the expected range for dimeric coiled-coils^28–30^. Furthermore, their interatomic distance distribution *P(r)* indicates maximum dimensions of 170 Å and 180 Å, respectively (**Figure 3B**), slightly longer than the length (160 Å) of the cNTD in the crystal. Fitting of the cNTD crystal structure to the SAXS data gave moderate fits, with χ^2^ values of 6.94 (WT) and 3.89 (Mut), which were improved by flexible modelling of disordered residues at the termini of the crystallographic model (χ^2^ of 5.08 and 2.75, respectively) (**Figure 3A** and **Supplementary** t**able 2**). Thus, the wild-type and mutant CtIP-cNTD proteins in solution contain subtle structural differences relative to the crystal structure of the mutant CtIP-cNTD.

**Figure 3.**
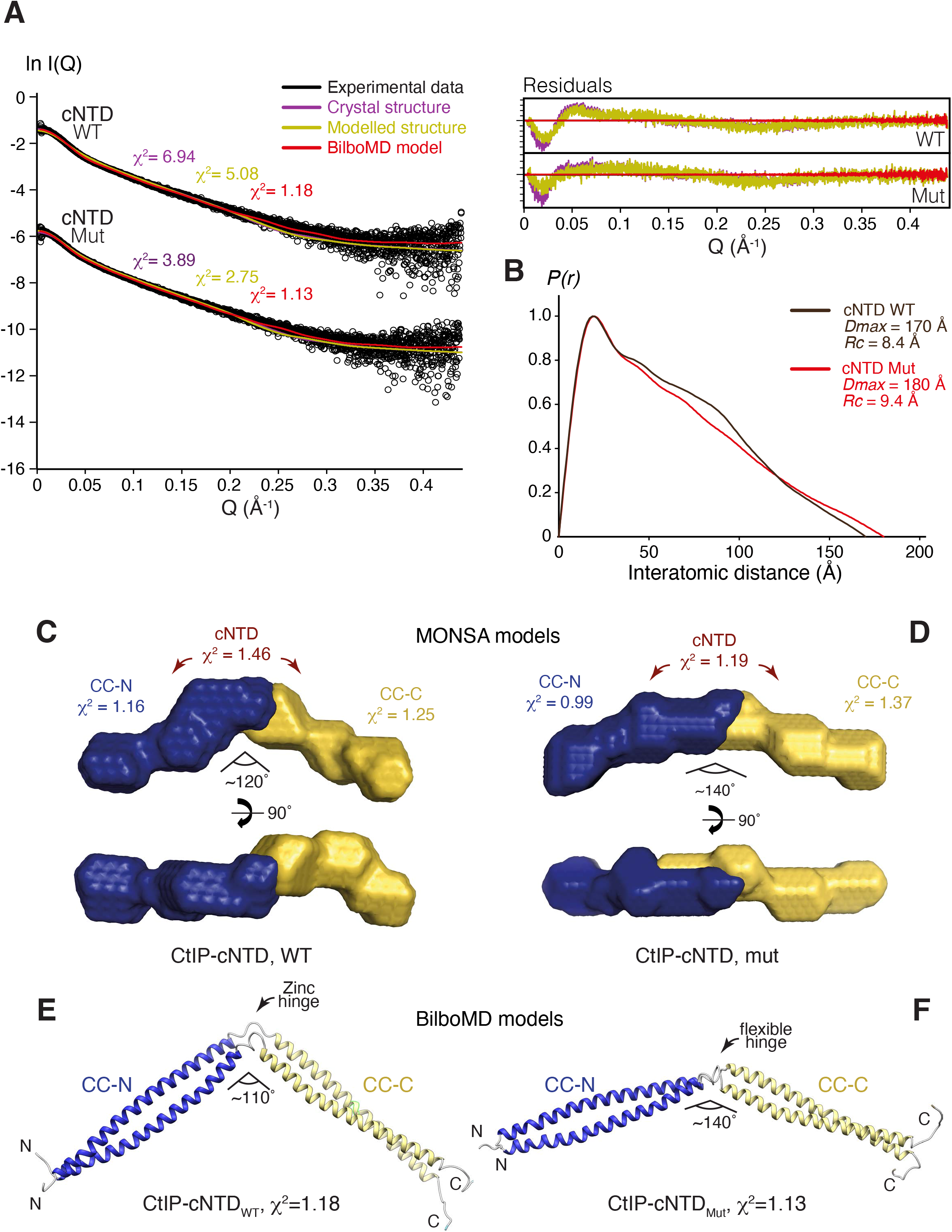
CtIP-cNTD contains a CxxC zinc-hinge. **A** SEC-SAXS analysis of wild-type and mutant cNTD. SAXS scattering data (black) are overlaid with the theoretical scattering curves of the crystal structure (purple), the crystal structure remodelled at its C-terminus (yellow) and BilboMD molecular dynamics models (red). Residuals for each fit are shown (inset). **B** Interatomic distance distributions of wild-type (green) and double-mutant (red) cNTD, showing maximum dimensions of 170 Å, 180 Å, respectively. **C** and **D** Multi-phase SAXS *ab initio* (MONSA) models of wild-type and mutant cNTD, respectively, coloured blue (CC-N) and yellow (CC-C). χ^2^ values for the entire envelope and individual phases are shown. **E** and **F** Molecular dynamics (BilboMD) models of wild-type and mutant cNTD, respectively, generated with CC-N and CC-C as rigid domains. Colour scheme as in C and D.

To gain insight into the solution conformation of the cNTD protein, we performed multi-phase SAXS *ab initio* modelling of wild-type and mutant CtIP-cNTD in MONSA^31^. The *ab initio* modelling demonstrated the end-on arrangement of the CC-N and CC-C regions, with a distinct obtuse angle at the junction between CC-N and CC-C, corresponding to the C89xxC92 site (**Figure 3C, D**). This angle is more marked for the wild-type (120°) than the mutant protein (140°), suggesting that zinc coordination induces a kink to the cNTD structure that is reduced upon loss of zinc binding.

In parallel, we performed SAXS-directed modelling through a molecular dynamics (MD) approach using BilboMD^32^, in which dimeric regions of amino-acids 31-91 and 93-145 were defined as rigid domains within the cNTD model, with flexibility at the C89xxC92 site and the unstructured termini. The resultant models closely fitted the wild-type and mutant SAXS data, with χ^2^ values of 1.18 and 1.13 (**Figure 3A**) and demonstrated clear angulations at their C89xxC92 sites that are more marked in the wild-type, with a reduction from 140° to 110° in the mutant (**Figure 3E, F**).

Both *ab initio* and MD modelling of the cNTD indicate that zinc coordination at the C89xxC92 site imposes a pronounced angle between the CC-N and CC-C segments of the cNTD, akin to a zinc hinge. The loss of zinc-binding in the mutant likely disrupts this hinge conformation, allowing the near-linear helical conformation observed in the crystal structure of the cNTD mutant.

### SAXS analysis of the CtIP-NTD tetramer

How does the zinc hinge affect the conformation of the CtIP tetramer? In agreement with the model of CtIP-NTD as an elongated dimer-of-dimers, SAXS analysis of CtIP-NTD (18 – 145) revealed a cross-sectional radius *R*_*c*_ of 9.4 Å and maximum dimension *P(r)* of 330 Å (**Figure 4A, B**, **Supplementary Figure 2A, B** and **Supplementary Table 2**), indicating a coiled-coil structure of almost twice the length of a single cNTD molecule. Furthermore, multi-phase SAXS *ab initio* modelling demonstrated the end-on arrangement of two angled cNTD molecules (**Figure 4C**), consistent with the proposed head-to-head ‘dimer of dimers’ assembly.

**Figure 4.**
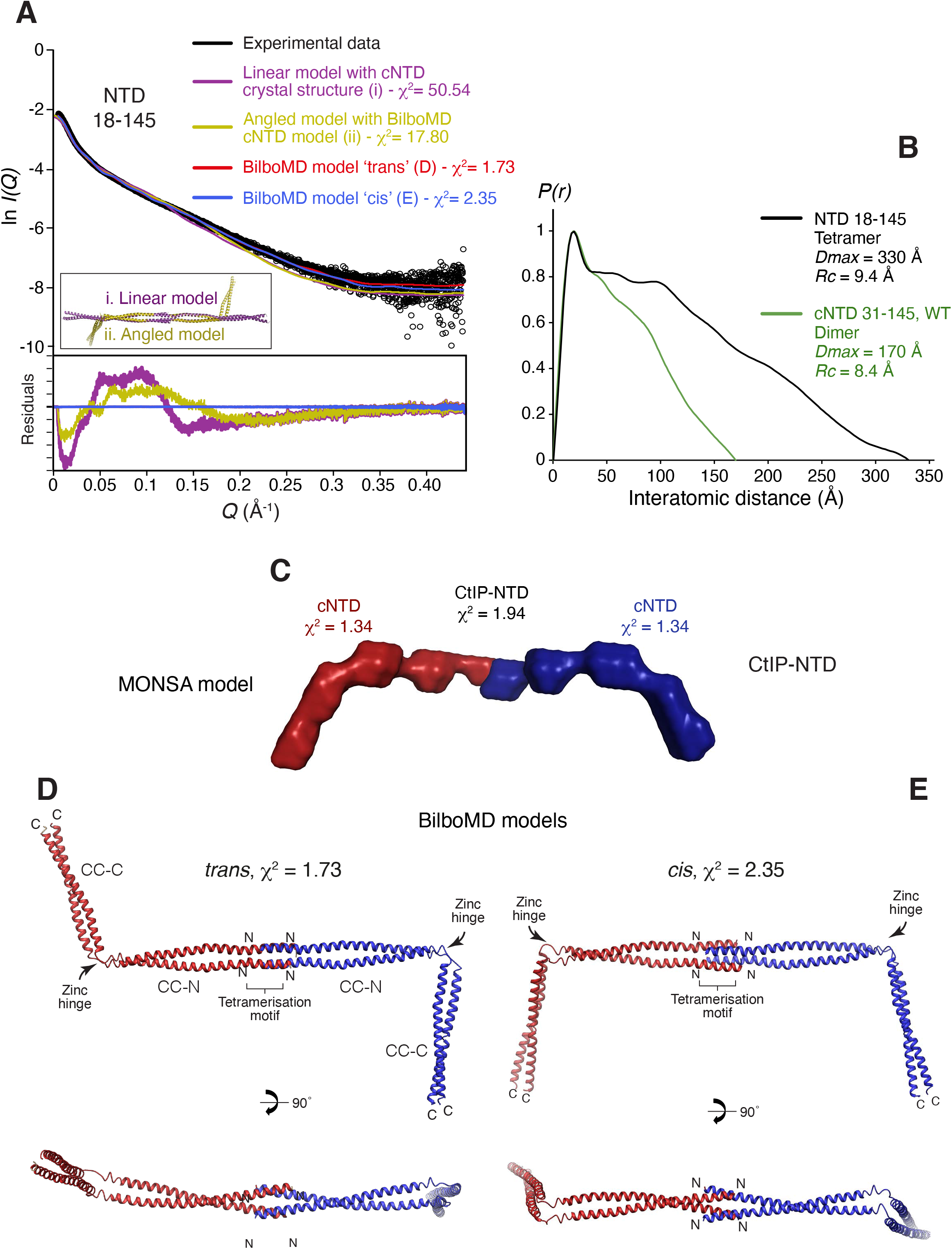
*Cis* or *trans* configurations of the CtIP-NTD tetramer. **A** SAXS analysis of CtIP-NTD (18 - 145). SAXS scattering data (black) overlaid with the theoretical scattering curves of linear (purple) and angled (yellow) models of the structure (shown in the inset), alongside BilboMD molecular dynamics models in *cis* (blue) and *trans* (red) configurations. Residuals for each fit are shown (inset). **B** Interatomic distance distribution of CtIP-NTD, showing maximum dimensions of 300 Å. The distance distribution of CtIP-cNTD is shown in green for comparison. **C** Multi-phase *ab initio* (MONSA) model of the CtIP-NTD tetramer, consisting of two cNTD dimers, in red and blue envelopes. **D, E** Molecular dynamics (BilboMD) models of CtIP-NTD, generated with the 18-88 tetramer and two 93-145 dimers as rigid bodies, in *trans* and *cis* conformations.

The nNTD and cNTD crystal structures share amino acids 31-52, allowing two cNTD molecules to be docked onto one nNTD structure, thus generating a three-dimensional model of the CtIP-NTD (**Supplementary Figure 2C**). Due to the presence of the kink caused by the zinc-binding motif in the middle of the cNTD, it was possible to generate CtIP-NTD models in either *cis* or *trans* configurations, with opposing cNTD molecules bending in the same or opposing directions. Whilst linear CtIP-NTD models generated from the cNTD crystal structure fitted poorly the experimental SAXS data (**Figure 4A**; χ^2^=50.54), fits were improved using angled CtIP-NTD models generated from the previously described cNTD molecular dynamics models (**Figure 4A**; χ^2^=17.80). On this basis, we performed SAXS-directed molecular dynamics modelling from *cis* and *trans* CtIP-NTD models in which the central 18-88 tetramer and two 93-145 dimers were specified as rigid bodies, with flexibility allowed within the C89xxC92 site. The resultant *cis* and *trans* models closely fitted the experimental data (χ^2^ values of 2.35 and 1.73, **Figure 4A**), and revealed clear angulations at the zinc-hinge, resulting in CC-C coiled-coils that are orientated in the same or opposing directions (**Figure 4D, E**).

This analysis suggests that the zinc hinge generates CtIP-NTD structures in which opposing cNTD molecules can adopt either *cis* or *trans* conformations, according to the relative direction of bending at the zinc hinges of the two cNTDs in the CtIP tetramer. We speculate that these alternative conformations of the CtIP-NTD could play a role in mediating the appropriate tetrameric CtIP conformation in the presence of different DNA-end geometries.

### Dynamic analysis of CtIP-cNTD conformation by Diffracted X-ray Tracking

To explore further the conformation of CtIP-cNTD, we performed a comparative analysis of wild-type and mutant CtIP-cNTD using Diffracted X-ray Tracking (DXT). DXT allows sensitive detection of dynamic changes in protein structure by tracking the X-ray diffraction spots of a gold nano-crystal covalently attached to the protein (**Figure 5A**)^33^. The motion of the diffraction spot generates a track across the detector that can provide dynamic information on intra-molecular tilting (theta (θ) angle) and twisting (chi (χ) angle) motions of the protein, as well as information on relative domain mobility when structural information is available^34^. For the DXT experiment, CtIP-cNTD was immobilised on the surface of the experimental support by maleimide coupling, which guaranteed a high degree of support derivatisation, and then labelled with gold nanocrystals. DXT experiments were performed using broad-band synchrotron radiation, and a 200 nano-second, time-resolved photon-counting detector. DXT traces were collected for wild-type and mutant CtIP-cNTD (**Supplementary figure 3A**).

**Figure 5.**
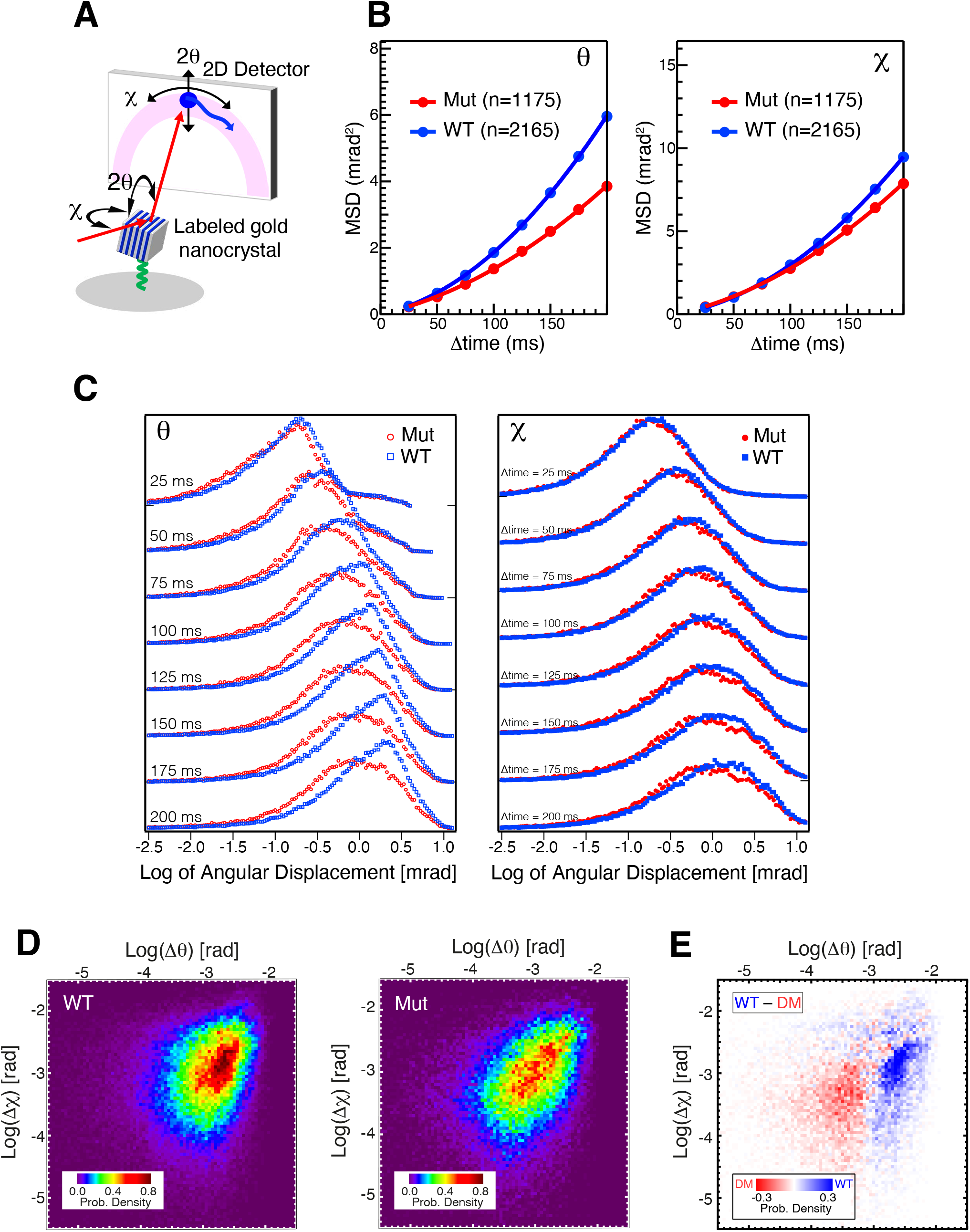
DXT analysis of CtIP-cNTD. **A** Schematic diagram of the setup for a DXT experiment. **B** Mean square displacement (MSD) curves for the θ and χ angles. ‘n’ is the number of diffraction spots tracked from the labeled gold nanocrystals targeted for DXT analysis. **C** The time-resolved distributions of angular displacements over a period of 200 ms for θ and χ. **D** Two-dimensional histogram representation of the angular displacement distribution for θ at 200 ms (panel C). **E** Two-dimensional difference histogram between the angular displacement distribution of θ for WT and Mut, shown in panel D.

Analysis of the DXT data for wild-type and mutant CtIP-cNTD shows a more dynamic behaviour of the wild-type protein relative to the mutant protein (**Figure 5B**, **Supplementary figures 3B** and **4**). In particular, the largest difference is in the theta angle distribution, which corresponds to a tilting motion of the structure (**Figure 5C-E**). In light of the known structural information for CtIP-cNTD, this difference in tilting dynamics can be interpreted as increased flexibility due to a hinge motion centred at the zinc-binding motif, present in the wild-type but not in the mutant protein. Thus, the DXT analysis supports the presence of a dynamic hinge motion enabled by the zinc-binding motif and expands our structural characterisation of the CtIP-cNTD as an angular coiled-coil structure with a dynamic joint at its centre.

## Discussion

In this paper, we present two important advances concerning the molecular architecture of human CtIP. We have shown that its N-terminal domain exists predominantly as a parallel dimeric coiled coil. We have further shown that the zinc-binding motif in the middle of the coiled coil dimer introduces a structural discontinuity that imparts both a pronounced angulation and a distinct dynamic behaviour to the protein shape. These results expand considerably our experimental knowledge of the molecular architecture of CtIP, when considered together with our previous elucidation of the structural basis for CtIP tetramerisation and the published evidence for a dumbbell shape of human CtIP^25^. Thus, CtIP forms an extended dimer of dimers, where two parallel dimers of coiled-coil polypeptides tetramerise by intermeshing the N-end of their chains, while projecting away from each other in opposing directions.

Such a tetrameric architecture confers upon CtIP the ability to bridge distant DNA sites, either on the same or on different chromosomes or sister chromatids, in agreement with its well-established functions in DNA DSB repair and meiosis (**Figure 6**). We note that the head-to-head dimer-of-dimers architecture of CtiP is strikingly reminiscent of the tetrameric architecture of synaptonemal complex protein SYCP1^30^. During prophase I of meiosis, SYCP1 directs the close pairing of meiotic chromosomes, crosslinking parental chromosomes by virtue of an extended self-assembly mechanism driven by dimer-of-dimers interactions of its helical N-termini. It is possible that CtIP might adopt a similar mechanism to hold together DNA ends in preparation for end resection. Thus, the use of an extended architecture based on head-to-head association of dimeric helical coiled-coils might be a general way to hold together distant nucleic-acid segments during metabolic processes that require recombination between homologous DNA molecules.

**Figure 6.**
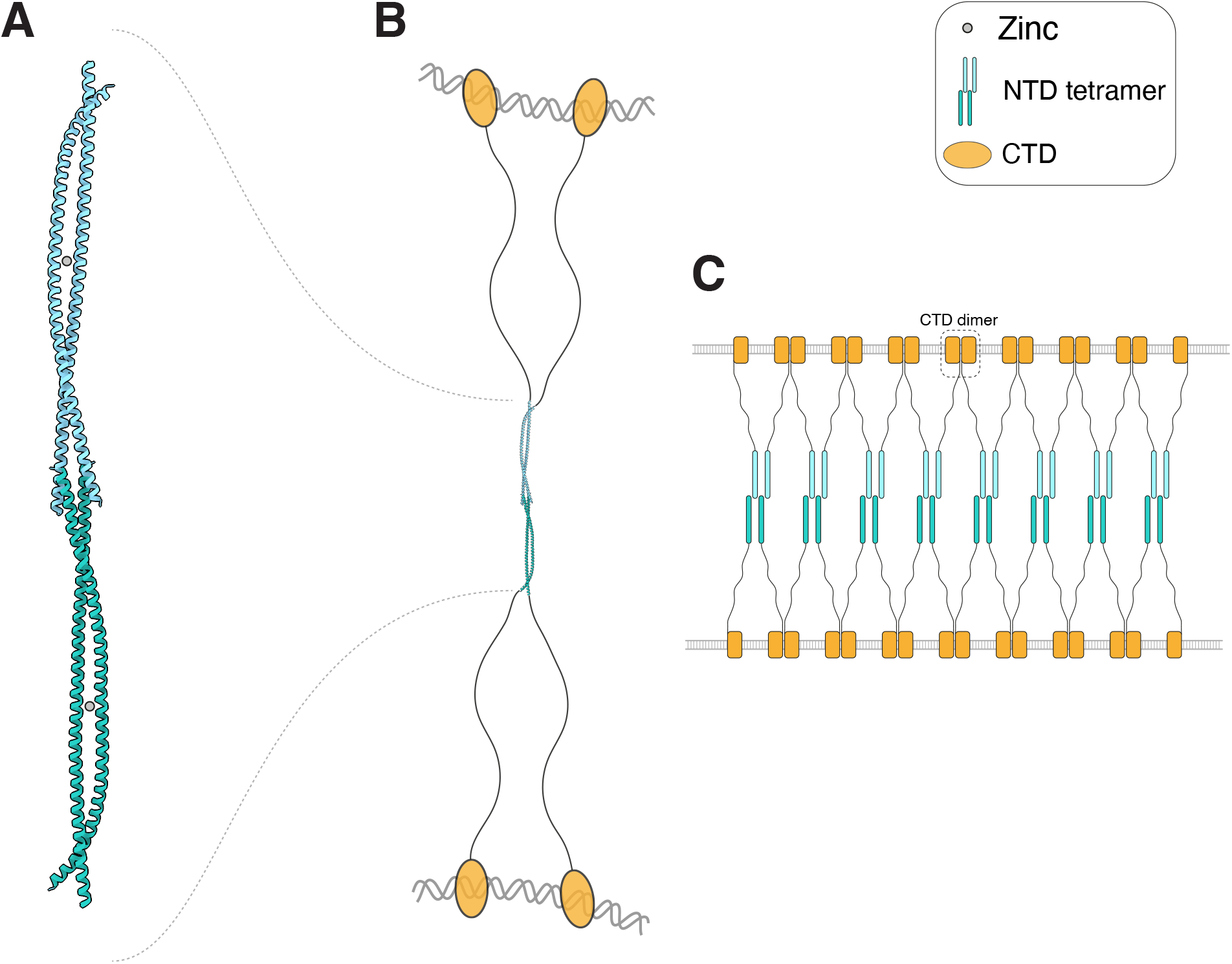
Model for the interaction of tetrameric CtIP with DNA. **A** Structure of the N-terminal CtIP sequence responsible for tetramer formation, obtained by merging the crystal structures for the tetramerisation motif and the dimeric coiled-coil region of CtIP. The protein chains are drawn as ribbons and the two dimers are coloured in cyan and green. **B** The architecture of full-length CtIP. Such dimer-of-dimers arrangement of CtIP chains would allow bridging of distant DNA sites via DNA-binding of the C-terminal domains (CTDs). The CTDs are shown as orange ovals, connected to the N-terminal tetrameric structure by intrinsically disordered regions. **C** A model for the possible multimerization of CtIP on DNA. In the model, juxtaposed dimers of dimers would connect via association of DNA-bound CTDs, potentially forming a protein network holding two DNA molecules together. The model is supported by recently published evidence of higher-order DNA-bound CtIP/Ctp1 structures^26,27^ and the ability of CTD to dimerise (CRM and LP, unpublished observation).

The role of the zinc-binding motif in such a mechanistic model of CtIP has remained unclear, as it is not required for CtIP-cNTD dimerization and no biochemical function, such as DNA binding, has been attributed to it; equally, no evidence for a putative function is available from cellular assays. Our data provides structural insight into the zinc-binding motif, by showing that its presence induces a marked angulation in the shape of the CtIP-cNTD. The consequence of such conformational effect on the coiled coil of the cNTD is that two different tetrameric architectures of CtIP can be envisioned; these can be purposefully named *cis* and *trans* according to whether the C-termini of the CtIP-NTD are found on the same side or opposite sides of the long two-fold symmetry axis of the tetramer.

The functional implications of the existence of two distinct arrangements for the CtIP-NTD are presently unclear. It is possible that the geometry of the DNA sites influences the protein conformation, so that a *cis* conformation might be selected in the case of a single DSB, whereas a *trans* conformation is adopted when holding onto pairs of sister chromatids. The fact that the majority of the CtIP protein beyond the NTD appears to be intrinsically unstructured highlights how such simple geometric models remain highly tentative. A deeper mechanistic understanding of CtIP function in DNA DSB repair will require high-resolution studies of the full-length protein, in isolation and bound within the end-resection complexes of Homologous Recombination.

## Acknowledgments

We would like to the beamline staff of Diamond Light Source (Nathan Cowieson, David Omar, Giulio Crevatin, William Nichols, Alan Greer, Nicola Tartoni) for help with SAXS and DXT data collection. This work was funded by Wellcome Trust investigator award 104641/Z/14/Z to L.P. O.R.D. is a Sir Henry Dale Fellow jointly funded by the Wellcome Trust and Royal Society (104158/Z/14/Z).

## Author contributions

CRM carried out all experimental work, assisted by NJR (protein preparation), JDM (CD experiments), RR (SAXS experiments); ORD performed the analysis of the SAXS data. CRM and YS performed the DXT experiments and YS, assisted by MK and HS, analysed the DXT data. LP conceived and supervised the project and wrote the manuscript with input from all authors.

Coordinates and structure factors for CtIP-cNTD have been deposited in the Protein Data Bank with access code 7BGF.

## Materials and Methods

### Recombinant-protein expression and purification

Sequences corresponding to human CtIP residues 18-145, 31-145, 31-88 and 91-145 including 31-145 double mutant C89A C92A, were cloned into pMAT11 vectors for expression as N-terminal His_6_-MBP fusion proteins with the addition of a Strep tag at the C-terminus. Constructs were expressed in BL21(DE3) Rosetta 2 cells (Novagen) in 2xYT medium, after induction with 0.5 mM IPTG for 16 h at 20°C. Fusion proteins were purified from clarified lysate by Ni-NTA (Qiagen) affinity chromatography. The His-MBP tag was removed by cleavage with TEV protease (Invitrogen). Further purification was achieved through capture of the cleaved protein by Strep-Tactin resin (IBA) and elution with 2.5mM *d*-desthiobiotin (Sigma). A final purification step was performed by size-exclusion chromatography using a HiLoad 16/60 Superdex 75 column (GE Healthcare) in 20 mM Tris HCl pH 8.0, 300 mM NaCl. Fractions containing pure CtIP protein were concentrated to 10 mg/ml and stored at −80 °C after flash freezing in liquid nitrogen.

### Circular dichroism spectroscopy

Far-UV CD spectroscopy data were collected on an Aviv 410 spectropolarimeter (Biophysics facility, Department of Biochemistry, University of Cambridge). CtIP samples were analysed at 0.2 mg/ml, in 5 mM Na_2_HPO_4_/NaH_2_PO_4_, pH 8.0 and 100 mM NaF, with a quartz cuvette with 1-mm path length. CD thermal denaturation data were recorded at 222 nm, at 2 °C intervals between 5 and 95 °C (1 °C/minute ramping rate, with 30-s incubation time). The data were fitted to a rearrangement of the Gibbs - Helmholtz relationship^35,36^ using pro Fit software (Quantum Soft).

### Size-exclusion chromatography multi-angle light scattering (SEC-MALS)

The absolute molecular masses of recombinant CtIP protein samples were determined by SEC-MALS. 100-μl protein samples (at approximately 2 mg/ml) were loaded onto a Superdex 200 or 75 10/300 GL Increase size-exclusion chromatography column (GE Healthcare) in 20 mM Tris HCl, pH 8.0, 300 mM NaCl, with or without 2 mM EDTA, at 0.5 ml/min with an ÄKTA Purifier (GE Healthcare). The column output was fed into a DAWN HELEOS II MALS detector (Wyatt Technology) followed by an Optilab T-rEX differential refractometer (Wyatt Technology). Light scattering and differential refractive index data were collected and analysed with ASTRA 6 software (Wyatt Technology). Molecular masses and estimated errors were calculated across individual eluted peaks by extrapolation from Zimm plots with a dn/dc value of 0.1850 ml/g. SEC-MALS data are presented with light scattering (LS) and differential refractive index (dRI) plotted alongside fitted molecular masses (Mr).

### Protein crystallization and X-ray structure solution

Initial crystallisation hits for the 31-145 DM CtIP protein were observed with condition F6 of the Morpheus^®^ commercial screen MD-47 (Molecular Dimensions). Crystals were improved by hanging-drop vapor diffusion and micro-seeding, mixing the protein sample at 7.5mg/ml with crystallisation buffer and micro seed sample at a ratio of 10:19:1. X-ray diffraction data were collected at wavelength 0.976251Å, 100K as 990 contiguous frames with 0.02 second exposure and 0.1° oscillation on a Pilatus3_6M detector at beamline ID30B of the ESRF Grenoble, France. The diffraction images were indexed and intensities extracted and scaled in XDS^37^ and further scaled and analysed using Aimless^38^. Crystals were assigned to the trigonal P3_2_ space group with unit cell dimensions: a=b=86.6Å, c=42.6Å, α=β=90, γ=120, with one protein dimer in the asymmetric unit. The crystal structure was solved by Molecular Replacement (MR) using Phaser^39^. Two helical coiled-coil models spanning amino acids 56-83 and 99-144 of human CtIP, corresponding to the regions of CtIP NTD strongly predicted to form coiled-coil structures, were generated in CBuilder2.0^40^ and used as search templates. Phaser successfully placed both models in the asymmetric unit, with no model clashes and clear indication of extra electron density for the missing parts of the crystallographic model. An initial poly-alanine model was refined in Phenix^41^, adding side chains and extending the model to boundaries of the interpretable density maps. Refinement in Phenix was interspersed with manual rebuilding in Coot ^42^, to improve fitting to the electron density map and the stereochemistry of the model. The refined crystallographic model of the human CtIP NTD comprise amino acids 31 to 136 for both chains, with R_work_/R_free_ = 0.2528/2235, no Ramachandran outliers and an overall Molprobity score of 1.05 ^43^.

### Size-exclusion chromatography small-angle X-ray scattering (SEC-SAXS)

SEC-SAXS experiments were performed at the bioSAXS beamline B21 (Diamond Light Source synchrotron, UK). Protein samples at concentrations in the range 3 to 10 mg/ml were analysed using a Superdex 200 Increase 3.2/300 2.4 ml column in 20 mM Tris HCl 8.0, 300 mM NaCl at 0.05 ml/min in an Agilent 1200 HPLC system. The column outflow passed through the experimental cell, where SAXS data were recorded at 12.4 keV, detector distance 4.014 m, in 3.0 s frames. ScÅtter 3.0 (http://www.bioisis.net) was used to subtract, average and carry out Guinier analysis for the *Rg* and cross-sectional *Rg* (*Rc*), and *P(r)* distributions were fitted using PRIMUS^44^. Multi-phase SAXS *ab initio* modelling was performed using MONSA^31^; rigid-body and flexible tail modelling was performed using CORAL^45^. Molecular dynamics modelling against experimental SAXS data was performed using BilboMD^32^ (https://bl1231.als.lbl.gov/bilbomd), with regions 18-88 or 31-88 and 93-145 specified as rigid bodies. Crystal structures and models were fitted to experimental data using CRYSOL^46^.

### Structural modelling

Structural modelling and visualisation were performed using Coot^42^ and PyMOL Molecular Graphics System, Version 2.3 Schrödinger, LLC. The CtIP-cNTD C-terminus was modelled as an ideal coiled-coil using CCBuilder 2.0^40^ (http://coiledcoils.chm.bris.ac.uk/ccbuilder2/builder), docked onto the cNTD crystal structure and used to replace the deviating terminus of chain A (amino-acids 122-138) and to extend chain B to amino acid 145. The tetrameric CtIP-NTD structure was modelled by docking two copies of the cNTD onto the nNTD crystal structure^21^ (PDB ID 4D2H) based on the shared structure of amino acids 31-52 (r.m.s. deviation = 0.413). Models were generated in *cis* or *trans* configurations by docking cNTD structures with their asymmetric bends in the same or opposing directions, respectively. Linear models were generated from the cNTD double-mutant crystal structure, whereas angled models were generated using the angled cNTD structure that was obtained from BilboMD molecular dynamics modelling against experimental CtIP-cNTD wild-type SAXS data.

### Diffracted X-ray Tracking

A Kapton (Polyimide film, DuPont) surface was pre-treated with UV radiation for 1/6 h to generate a negatively charged surface for poly-lysine (0.01% W/V) coating. To this, GMBS (N-(4-Maleimidobutyryloxy) succinimide (Dojindo Laboratories)) was bound to the poly-lysine coated surface. For sufficient CtIP-cNTD coverage, CtIP-cNTD was applied to the GMBS coated surface in excess and incubated at 4°C for 1/6 h. Gold nano crystals were pre-treated with 10-Carboxy-1-decanethiol (Dojindo Laboratories) to prevent aggregation and modify the gold surface for both methionine and cysteine coupling to CtIP-cNTD. This was subsequently incubated with the protein surface for 1/6h at 4°C followed by washing into the protein buffer and sealing within the apparatus for exposure to the X-ray beam.

DXT experiments were performed at the B16 beamline of the Diamond Light Source synchrotron (UK). The CtIP-cNTD samples were irradiated with X-rays with energy bandwidths ranging from 11 to 17 keV and with beam size of 0.5 mm diameter, and diffraction movies were recorded by a Tristan 1M detector with 16 timepix3 chips located 120 mm from the sample. All diffracted photon hits on the detector were recorded and time-resolved diffraction images (25 ms/f) were reconstructed with the software provided by the detector group at Diamond Light Source. An incident X-ray irradiated one position of the sample for 3 seconds, and the same measurement was repeated at 50 different sample position on each sample. The motions of the spots diffracted from the gold nanocrystals on the substrate’s surface were tracked by TrackPy (v0.4.2 https://doi.org/10.5281/zenodo.3492186), and the trajectories of the diffraction spots (Supplementary Fig. 3A) were analysed using custom software written within IGOR Pro (Wavemetrics, Lake Oswego, OR). The motionless diffracted spots, less than 1 mrad in theta direction, and the events on the edge pixels in each Timepix3 chip were excluded for the analysis.

