## Supplementary information for "Structural basis for the coiled-coil architecture of human CtIP"

**Supplementary table 1. Data collection and refinement statistics.**

| | CtIP-cNTD |
|---|---|
| *Data collection* | |
| Wavelength | 0.9763 |
| Resolution range | 37.5 - 2.802 (2.902 - 2.802) |
| Space group | P 32 |
| Unit cell | 86.598 86.598 42.601 90 90 120 |
| Total reflections | 24133 (2432) |
| Unique reflections | 8624 (878) |
| Multiplicity | 2.8 (2.8) |
| Completeness (%) | 97.95 (99.10) |
| Mean I/sigma(I) | 12.18 (1.29) |
| Wilson B-factor | 79.89 |
| R-merge | 0.05463 (0.7581) |
| R-meas | 0.06769 (0.9365) |
| R-pim | 0.03947 (0.5435) |
| CC1/2 | 0.999 (0.601) |
| *Refinement* | |
| Reflections used in refinement | 8612 (877) |
| Reflections used for R-free | 813 (88) |
| R-work | 0.2235 (0.3263) |
| R-free | 0.2528 (0.3781) |
| CC(work) | 0.415 (-0.024) |
| CC(free) | 0.528 (-0.024) |

| | |
|---|---|
| Number of non-hydrogen atoms | 1913 |
| macromolecules | 1877 |
| solvent | 36 |
| Protein residues | 220 |
| RMS(bonds) | 0.002 |
| RMS(angles) | 0.26 |
| Ramachandran favored (%) | 100.00 |
| Ramachandran allowed (%) | 0.00 |
| Ramachandran outliers (%) | 0.00 |
| Rotamer outliers (%) | 0.48 |
| Clashscore | 2.62 |
| Average B-factor | 101.61 |
| macromolecules | 102.02 |
| solvent | 80.05 |

Statistics for the highest-resolution shell are shown in parentheses.

$$R_{merge} = \frac{\sum_{hkl} \sum_{j} |I_{hkl,j} - \langle I_{hkl} \rangle|}{\sum_{hkl} \sum_{j} I_{hkl,j}}$$

$$R_{meas} = \frac{\sum_{hkl} \sqrt{\frac{n}{n-1}} \sum_{j=1}^{n} |I_{hkl,j} - \langle I_{hkl} \rangle|}{\sum_{hkl} \sum_{j} I_{hkl,j}}$$

$$R_{pim} = \frac{\sum_{hkl} \sqrt{\frac{1}{n-1}} \sum_{j=1}^{n} |I_{hkl,j} - \langle I_{hkl} \rangle|}{\sum_{hkl} \sum_{j} I_{hkl,j}}$$

$$R\ (Rf_{ree}) = \frac{\sum_{hkl} |F_{hkl}^{obs} - F_{hkl}^{calc}|}{\sum_{hkl} F_{hkl}^{obs}}$$

### Supplementary table 2. Summary of SAXS data fitting

| | **CtIP** NTD (18-145) | **CtIP** cNTD (31-145) Wild-type | **CtIP** cNTD (31-145) DM |
|---|---|---|---|
| **SEC-SAXS** | | | |
| *I(0)* ($cm^{-1}$) (from *P(r)*) | $1.20x10^{-1}$ | $6.00x10^{-2}$ | $5.00x10^{-2}$ |
| *I(0)* ($cm^{-1}$) (from Guinier analysis*)* | $1.22x10^{-1}$ | $5.53x10^{-2}$ | $4.91x10^{-2}$ |
| *Rg* (Å) (from *P(r)*) | 95.0 | 50.1 | 51.5 |
| *Rg* (Å) (from Guinier analysis) | 89.9 | 47.8 | 47.1 |
| *Rc* (Å) | 9.4 | 8.4 | 9.4 |
| *Dmax* (Å) | 330 | 170 | 180 |
| Porod volume ($Å^3$) | 118477 | 51285 | 52417 |
| Molecular weight (from Porod volume) (kDa) | 69.7 | 30.2 | 30.8 |
| Volume of correlation ($V_c$) ($Å^2$) | 762.1 | 379.6 | 373.3 |
| Molecular weight (from $V_c$) (kDa) | 52.5 | 24.5 | 24.0 |
| Crystal structure fit ($\chi^2$) | N/A | 6.94 | 3.89 |
| Modelled structure fit ($\chi^2$) | 50.54 (linear/trans)<br>48.87 (linear/cis)<br>17.80 (angled/trans)<br>16.18 (angled/cis) | 5.08 | 2.75 |
| CORAL model fit ($\chi^2$) | N/A | 3.40 | 2.23 |
| BilboMD model fit ($\chi^2$) | 1.73 (trans)<br>2.35 (cis) | 1.18 | 1.13 |
| MONSA multi-phase *ab initio* model fit ($\chi^2$) | 1.94 | 1.46<br>1.34 | 1.19 |

**Supplementary figure 1.** SAXS analysis of WT and Mut CtIP-cNTD. Guinier analysis was used to determine the radius of gyration ($R_g$; panel **A**) and cross section ($R_c$; panel **B**) for WT cNTD and Mut cNTD. Linear fits are shown in red, with the fitted data range highlighted in black and demarcated by dashed lines.

**Supplementary figure 2.** SAXS analysis of tetrameric CtIP-NTD. SAXS Guinier analysis to determine the radius of gyration (*Rg*) (**A**) and cross section (Rc) (**B**). Linear fits are shown in red, with the fitted data range highlighted in black and demarcated by dashed lines. **C** Model of the CtIP-NTD structure generated by docking two copies of the cNTD crystal structure onto the nNTD crystal structure (PDB ID 4D2H).

**Supplementary figure 3.** DXT traces and Gaussian fitting of the DXT data for CtIP-cNTD. **A** Trajectories of diffraction spots on the TRISTAN-1M detector from gold nanocrystals immobilized on WT (upper) and Mut (bottom) CtIP-cNTD proteins. The detector is made up of 16 (8 x 2) timepix3 chips and chip-to-chip gaps were observed in these images. **B** The distributions of angular displacements during 200 ms for the $\theta$ and $\chi$ angles. All motion distributions were fitted with a four-Gaussian distribution (see Supplementary Figure 4). In all distributions, the WT protein had a larger motion distribution than the Mut protein.

**Supplementary figure 4.** Maximum likelihood estimates of optimal number of components for gaussian fitting of the DXT data. By this fitting method, it was evaluated that the motion distribution measured by DXT was best represented by four Gaussians.

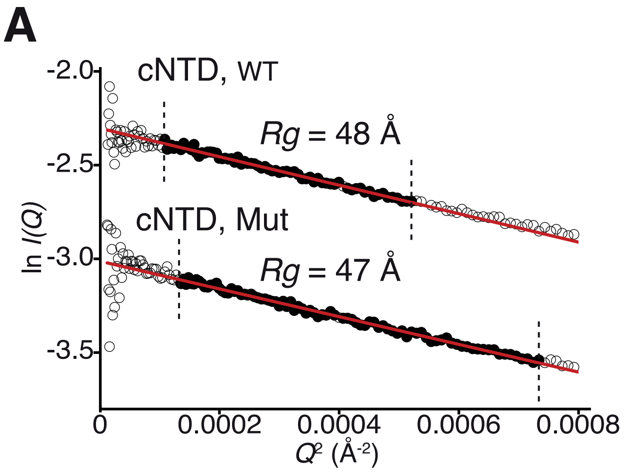


**Supplementary figure 1**

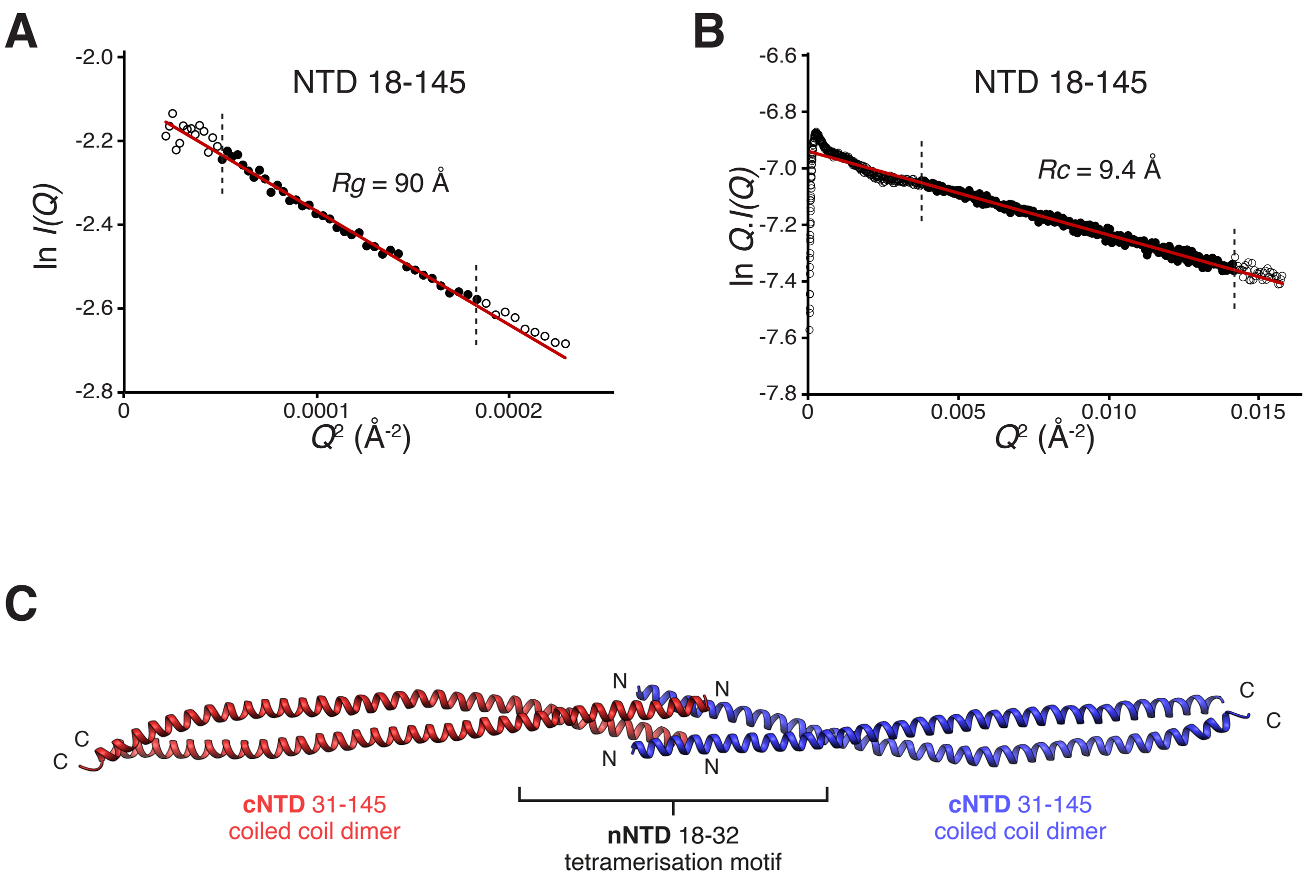


**Supplementary figure 2**

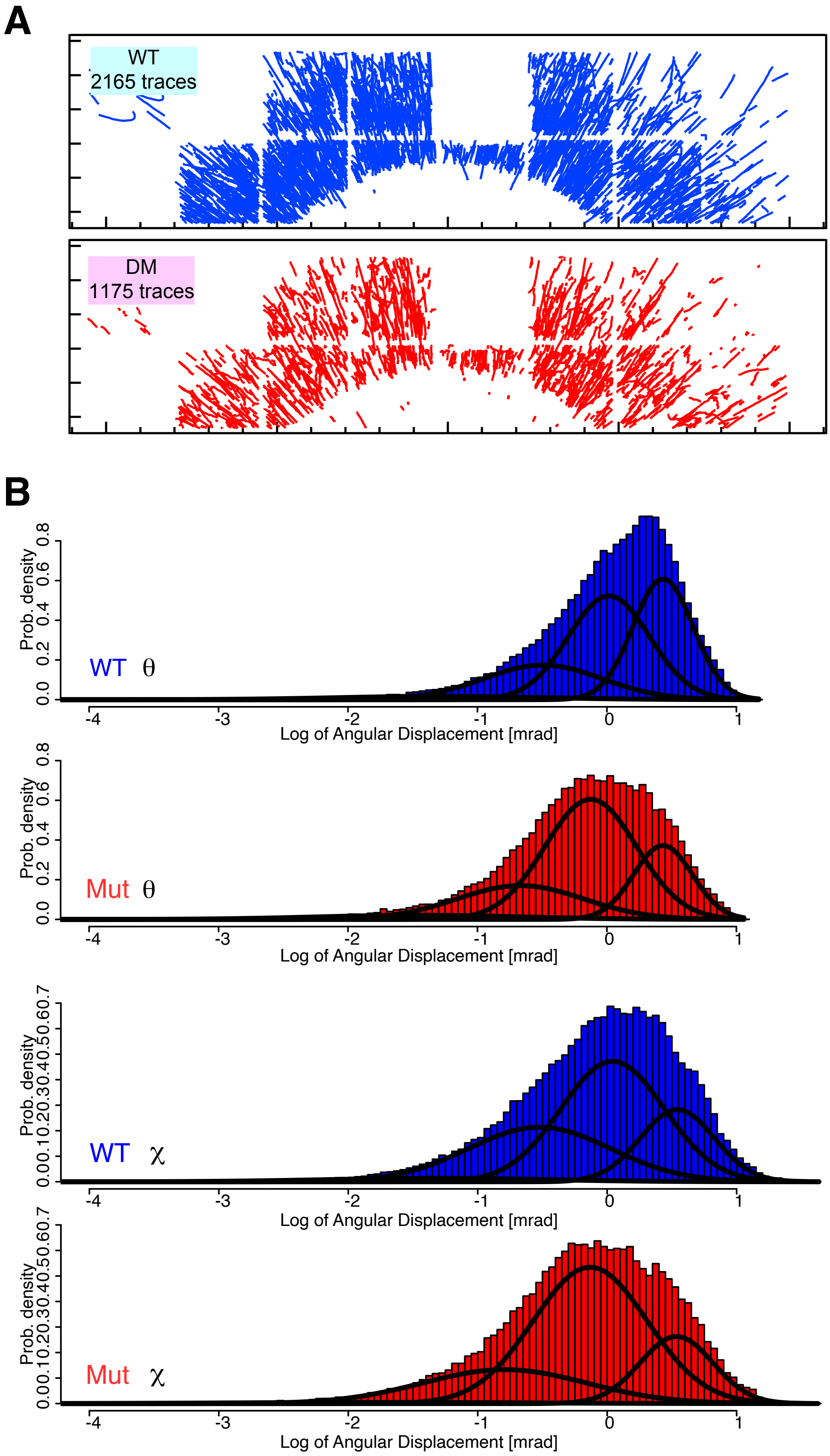


**Supplementary figure 3**

Maximum Likelihood Estimation with Expectation–Maximization algorithm
– How many Gaussians? (θ, ΔT = 200ms, WT vs DM)

Comparison of log-maximum likelihood

| No of Gaussians | WT | DM |
|---|---|---|
| 1 | -67139.18 | -41705.18 |
| 2 | -63266.67 | -40328.11 |
| 3 | -61930.37 | -39762.34 |
| 4 | -61733.65 | -39653.08 |
| 5 | -61614.49 | -39628.64 |
| 6 | -61561.6 | -39609.79 |
| 7 | -61559.42 | -39602.6 |
| 8 | -61593.72 | -39577.16 |

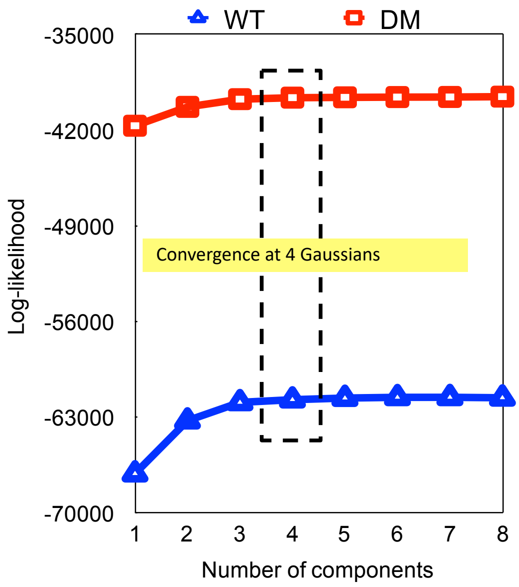


Maximum Likelihood Estimation with Expectation–Maximization algorithm
– How many Gaussians? (χ, ΔT = 200ms, WT vs DM)

Comparison of log-maximum likelihood

| No of Gaussians | WT | DM |
|---|---|---|
| 1 | -82525.75 | -47437.3 |
| 2 | -80281.63 | -46519.02 |
| 3 | -79673.7 | -46160.51 |
| 4 | -79564.95 | -46049.84 |
| 5 | -79541.24 | -46044.08 |
| 6 | -79540.74 | -46048.45 |
| 7 | -79542.37 | -46041.99 |
| 8 | -79515.87 | -46046.19 |

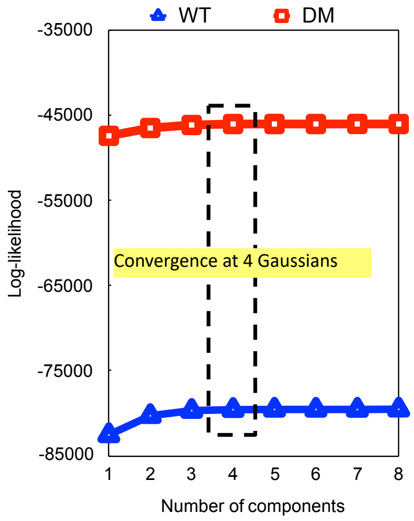


**Supplementary figure 4**
